# *Hamelia patens*-Derived Red-Emitting Carbon Quantum Dots: Surface-State Luminescence, Antioxidant Potency, and In Vitro Bioimaging

**DOI:** 10.64898/2026.05.10.724069

**Authors:** Satyam Bhalerao, Jugal Patil, Abdul Mansuri, Sweny Jain, Sanjay Kosara, Geethu Prakash, Ashutosh Kumar, Dhiraj Bhatia

## Abstract

Red-emitting carbon quantum dots (HP-CQDs) were synthesised for the first time from aqueous leaf extracts of Hamelia patens through single-step, reagent-free microwave-assisted carbonisation (750 W). The resulting nanoparticles displayed a narrow hydrodynamic size distribution centred at 3.9 nm, consistent with atomic force microscopy measurements showing a maximum height of 2.81 nm. Under 400 nm excitation, the CQDs exhibited a characteristic red emission maximum at 675 nm, representing a rare example of long-wavelength-emitting green CQDs derived from plant biomass. UV-Vis absorption bands at 224 and 256 nm were assigned to π-π* transitions of aromatic carbon domains and n-π* transitions associated with carbonyl-containing surface groups, respectively. X-ray photoelectron spectroscopy (XPS) indicated a carbon-rich composition (C: 67.24%, O: 31.25%, N: 1.52%) with prominent C-O (42.67%) and C-C/C=C (42.64%) contributions. ATR-FTIR further confirmed the retention of hydroxyl, ether, and aliphatic functionalities following carbonisation. The excitation-wavelength-independent emission peak position implicates discrete surface molecular states rather than a heterogeneous distribution of emitters. HP-CQDs exhibit potent DPPH radical scavenging activity (ICOO = 141.8 µg mLO¹), comparable to ascorbic acid (ICOO = 114.8 µg mLO¹), and maintain >95% cell viability in both HeLa and RPE-1 cells up to 250 µg mLO¹. Confocal microscopy demonstrates concentration-dependent cytoplasmic accumulation and selective perinuclear localization at 300 µg mLO¹. In vivo biodistribution in zebrafish larvae confirms systemic uptake with statistically significant fluorescence enhancement at 500 µg mLO¹ (p < 0.01), establishing HP-CQDs as biocompatible red-fluorescent probes with dual imaging-antioxidant functionality.

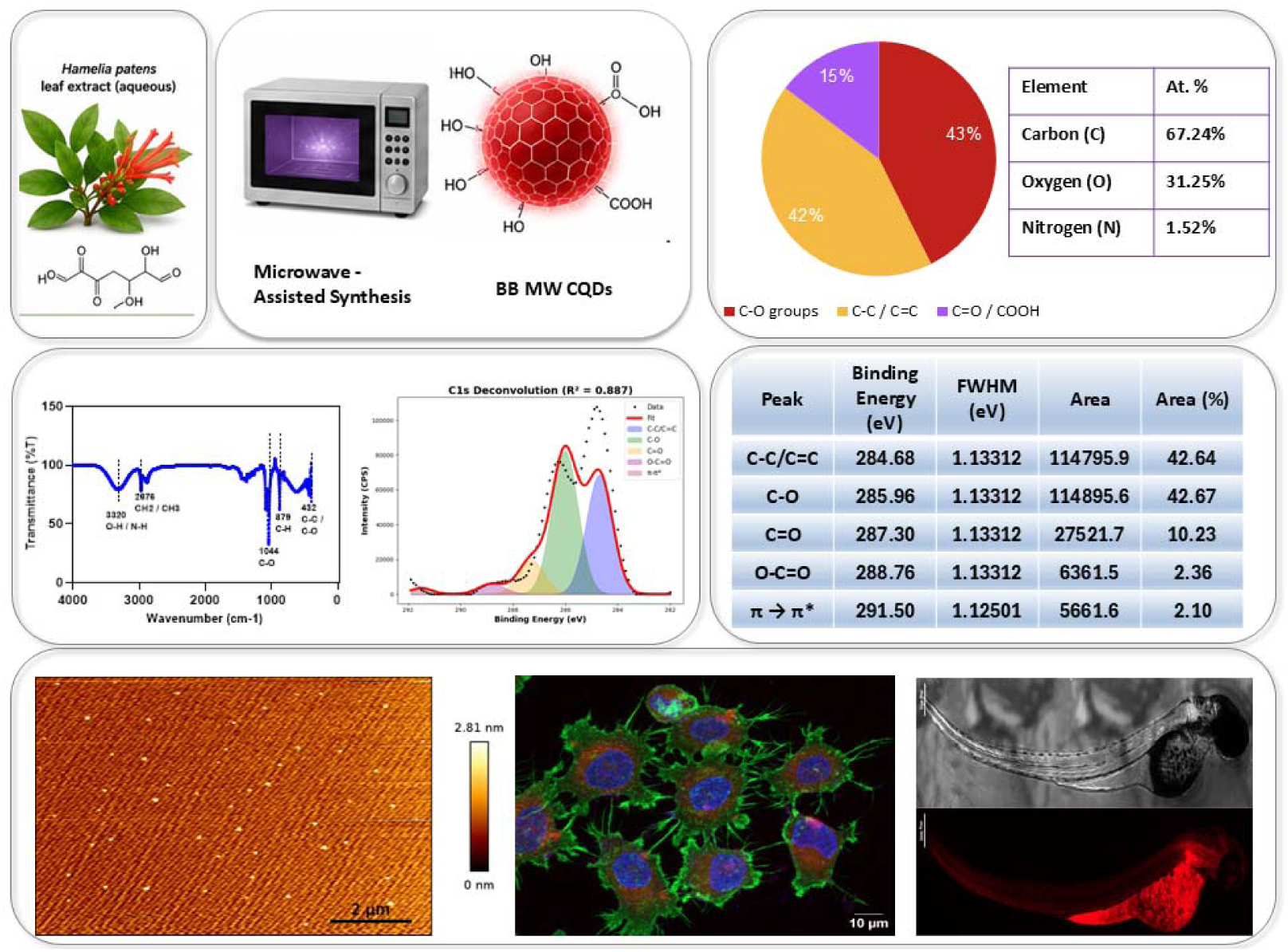

## 1. Introduction

Fluorescent nanomaterials have transformed the landscape of biomedical imaging, theranostics, and cellular diagnostics, largely owing to their photostability and size-tunable optical properties^1–4^. Among these, semiconductor quantum dots (QDs) based on CdSe, InP, and related heavy-metal chalcogenides exhibit excellent brightness and colour tunability, yet their inherent cytotoxicity and difficult surface passivation continue to restrict clinical translation^5^. Carbon quantum dots (CQDs) were first identified in 2004. They were observed as fluorescent by-products during the purification of single-walled carbon nanotubes. Since then, CQDs have emerged as promising nanomaterials. They are generally non-toxic, water-dispersible, and photostable^6,7^. CQDs can be synthesised from a wide range of carbon precursors^8–10,10,11^. Multiple fabrication strategies have been reported^12^. These include hydrothermal treatment, pyrolysis, electrochemical cutting, and microwave irradiation^3,13^.

Despite substantial progress, most plant-derived CQDs reported in the literature exhibit emission in the blue-to-green spectral window (420-550 nm). This behavior is primarily attributed to a limited conjugation length within the graphitic core. It is further influenced by the dominance of carboxyl and hydroxyl surface trap states. These states preferentially relax at shorter wavelengths. Red-emitting CQDs (emission > 600 nm) remain comparatively rare^14,15^. When achieved, they generally rely on additional modification strategies. These include nitrogen or sulfur co-doping using synthetic reagents such as urea, ethylenediamine, or thiourea^16^. In many cases, prolonged hydrothermal treatment (6-24 h) is required^16^. Post-synthetic surface passivation is also frequently employed. Such approaches compromise the ecological credentials and scalability that originally motivated green synthesis^17^. The photon-energy advantage of red emission reduced tissue autofluorescence, deeper penetration, and compatibility with far-red detector channels makes this spectral window highly coveted for in vitro and in vivo imaging^7,15,18,19^.

Hamelia patens (family Rubiaceae), commonly referred to as firebush or scarlet bush, is a pantropical shrub. Its leaves are rich in diverse phytochemicals. These include polyphenols, flavonoids such as rutin and quercetin, alkaloids, and chlorogenic acid derivatives. These biomolecules possess extended aromatic frameworks. They also contain multiple hydroxyl functionalities and conjugated carbonyl groups^20^. Under microwave-driven dehydration and carbonisation, such features are conducive to forming graphitised carbon cores. The resulting structures can support extended π-conjugation. This structural evolution is favourable for long-wavelength emission. To date, the use of Hamelia patens as a carbon precursor for CQD synthesis remains unexplored. The mechanistic relationship between its distinctive phytochemical composition and the emergence of red fluorescence has not been systematically investigated.

Microwave-assisted synthesis enables rapid and volumetric heating. This process ensures uniform carbonisation of the precursor. It avoids the diffusion limitations associated with conventional thermal pyrolysis. As a result, CQDs with narrower size distributions and more reproducible optical properties are obtained^21^. Notably, this can be achieved within minutes rather than hours. The integration of a renewable plant-derived precursor with microwave-driven carbonisation eliminates the need for auxiliary chemical reagents. This strategy aligns with principles of green nanotechnology. It also enhances the translational potential of CQDs for bioimaging applications.

In this work, we report the microwave-assisted synthesis of red-emitting HP-CQDs derived from Hamelia patens leaf extract. The synthesis was performed at 750 W in a water/ethanol medium. The product was subsequently processed through syringe filtration, rotary evaporation, and lyophilisation. Comprehensive characterisation was conducted to establish physicochemical properties. This included UV-Vis absorption, photoluminescence (PL) spectroscopy, and excitation–emission matrix (EEM) fluorescence. Structural and surface analyses were performed using ATR-FTIR and XPS, including high-resolution C 1s deconvolution. Colloidal and morphological properties were evaluated using DLS, zeta potential measurements, and atomic force microscopy (AFM).

Biological assessment was carried out to determine functional applicability^22^. Cytotoxicity was evaluated using MTT assays on HeLa and RPE-1 cell lines. Antioxidant activity was assessed using the DPPH radical scavenging assay, benchmarked against ascorbic acid. Cellular uptake and subcellular localisation were examined using confocal fluorescence microscopy. In vivo biodistribution was investigated using zebrafish larvae models.

Collectively, the results indicate that the observed red emission of HP-CQDs originates from surface-confined molecular states. These states are associated with aromatic O–C=O and C=O functional groups generated during microwave-induced carbonisation of a polyphenol-rich precursor. The resulting CQDs exhibit biocompatibility, antioxidant activity, and efficient cellular internalisation. These properties support their potential as next-generation fluorescent probes.

## 2. Results and Discussion

### 2.1 Optical Properties

#### 2.1.1 UV-Vis Absorption

The UV-Vis absorption spectrum of HP-CQDs in aqueous dispersion exhibits two well-resolved maxima at 224 nm and 256 nm (**Figure 1B**). The higher-energy band at 224 nm is attributed to π–π* transitions within aromatic sp²-hybridised carbon domains of the graphitic core, a feature characteristic of carbon nanoparticles derived from polycyclic aromatic precursors^23^. The band at 256 nm is assigned to n–π* transitions associated with surface C-O and C–O–C functionalities, consistent with the significant oxygenated carbon content confirmed by XPS (vide infra)^24^. Notably, the ground-state spectrum is featureless beyond 400 nm, yet HP-CQDs emit red photoluminescence under 365 nm UV excitation, manifesting as vivid pink emission in the fluorescence photograph (**Figure 1C**). This apparent decoupling of absorption and emission indicates that the emissive state is not populated through a direct far-visible ground-state transition; rather, emission arises via excited-state relaxation through surface-localised energy levels situated below the particle’s optically transparent window^25^. This behaviour is distinct from that of conventional molecular dyes, which absorb directly in the red, and is instead consistent with the excited-state surface-trap emission model described for oxygen-rich CQDs^26^.

**Figure 1.**
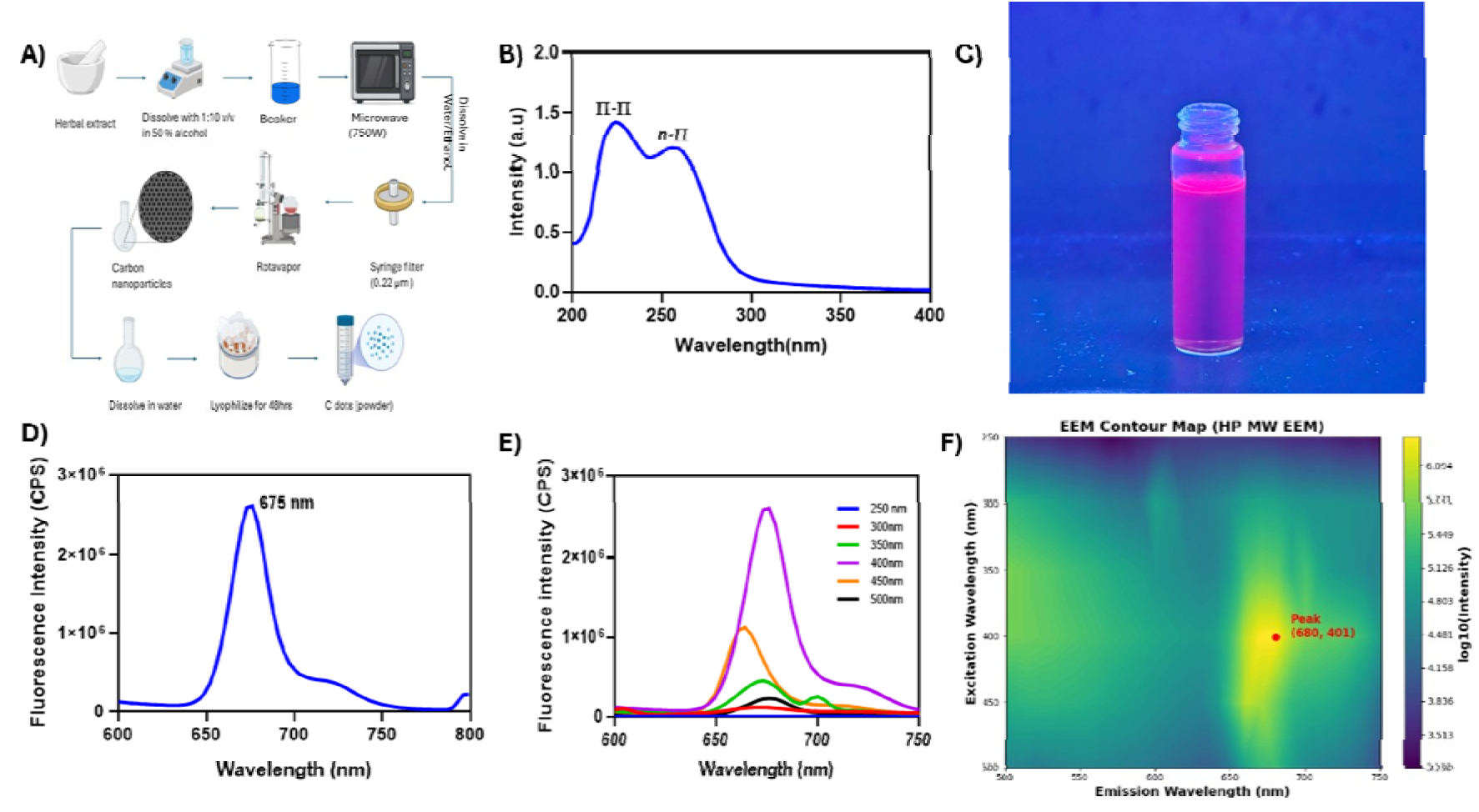
(A) Schematic of the microwave-assisted synthesis route for HP-CQDs from Hamelia patens leaf extract. (B) UV–Vis absorption spectrum showing π–π* (224 nm) and n–π* (256 nm) transitions. (C) Photographic image of HP-CQD colloidal dispersion under 365 nm UV irradiation, demonstrating red fluorescence. (D) Steady-state PL spectrum under 400 nm excitation, showing emission maximum at 675 nm. (E) Excitation-dependent PL spectra (λex = 250–500 nm). (F) Excitation-emission matrix (EEM) contour map with primary fluorescence peak annotated at (λem = 680 nm, λex = 401 nm).

#### 2.1.2 Photoluminescence Behaviour and Emission Characteristics

The steady-state photoluminescence spectrum of HP-CQDs, recorded under 400 nm excitation, exhibits a single and symmetric emission band centred at 675 nm. The fluorescence intensity exceeds 2.5 × 10LJ counts per second (CPS). This confirms the material as a genuine red-emitting nanostructure (**Figure 1D**). Upon illumination with a 365 nm UV lamp, the dispersion displays a bright pink-red emission visible to the naked eye. This provides direct qualitative evidence of long-wavelength emission from a plant-derived CQD system. The high emission intensity and pronounced spectral symmetry indicate a relatively uniform population of emissive states. This behaviour contrasts with many CQD systems that exhibit broad and asymmetric emission profiles. Such heterogeneity is typically associated with wide size distributions and multiple emissive centres.

Excitation-dependent PL spectra recorded over the 250–500 nm range (**Figure 1E**) reveal a mechanistically informative trend. The emission maximum remains fixed at ∼675-680 nm across all excitation wavelengths. In contrast, the emission intensity shows a pronounced maximum at 400 nm. It decreases at both shorter (250-350 nm) and longer (450-500 nm) excitation wavelengths. This behaviour differs from the classical excitation-dependent red-shift observed in heterogeneous CQDs^27^. In such systems, the emission peak typically shifts to longer wavelengths with increasing excitation wavelength. That effect is commonly attributed to multiple emissive states arising from structural and surface heterogeneity. In the present case, the invariant emission wavelength combined with excitation-dependent intensity indicates a single dominant emissive state. This state is most efficiently populated when the excitation matches the n–π* absorption transition near 400 nm^27^. This interpretation is further supported by the excitation–emission matrix (EEM) contour map (**Figure 1F**). The map shows a single high-intensity fluorescence feature at coordinates λ_em ≈ 680 nm and λ_ex ≈ 401 nm. No secondary emission centres are observed. This confirms the spectral purity of the emitting species^27^.

The mechanistic interpretation of this behaviour is as follows. Microwave-assisted carbonisation of Hamelia patens phytochemicals generates a graphitic CQD core with a surface densely functionalised with carbonyl, ether, and hydroxyl groups. These surface oxygen functionalities introduce discrete molecular energy levels below the carbon core band gap. When excited near 400 nm (matching the n-π* transition), the system undergoes rapid internal conversion to the lowest-energy surface trap state, from which radiative relaxation occurs at 675 nm. Excitation at shorter wavelengths (250-300 nm, π-π* region) can also populate this surface state via cascade relaxation from the core-excited state, but with lower efficiency due to competing non-radiative pathways accessible from the higher-energy core states. At longer excitation wavelengths (450-500 nm), the photon energy is insufficient to directly populate the primary absorbing chromophore, resulting in low emission intensity^19^. This surface-state mechanism, rather than quantum confinement, rationalises the red emission from particles as small as 3.9 nm, where quantum-confinement-driven red shifts in graphitic carbon would require significantly larger core diameters. The surface O-C=O moiety (2.36% by XPS, 288.76 eV) is particularly implicated as it possesses the extended n-π* absorption profile and the requisite energy gap for red emission, consistent with computational predictions for carboxy-functionalised graphene fragments^28^.

### 2.2 Surface Chemistry and Chemical State Analysis

#### 2.2.1 ATR-FTIR Spectroscopy

ATR-FTIR spectra of HP-CQDs (microwave-synthesised) and the parent Hamelia patens aqueous extract are compared in **Figure 2**. This comparison highlights the structural transformations induced during carbonisation. The extract spectrum (**Figure 2B**) is dominated by a broad and intense absorption band at 3389 cmO¹. This feature is assigned to O-H and N-H stretching vibrations arising from polyphenolic hydroxyl and amine groups in the native phytochemicals^11^. A second prominent band appears at 1644 cmO¹. This corresponds to overlapping C=O stretching of flavonoid carbonyl groups and C=C stretching of aromatic rings. The spectrum is relatively simple overall. Notably, aliphatic C-H stretching features are absent. This indicates that the extract is largely composed of aromatic and polar constituents.

**Figure 2.**
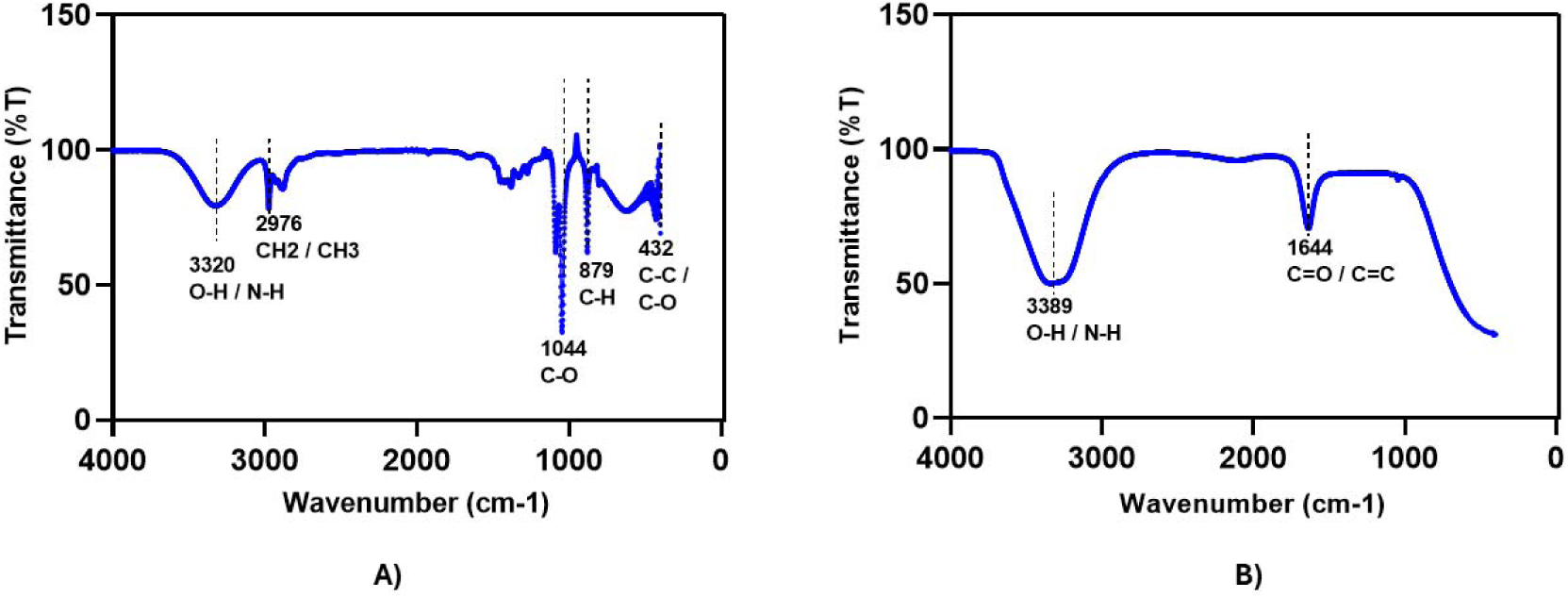
ATR-FTIR spectra of (A) HP-CQDs (microwave-synthesised) and (B) Hamelia patens leaf extract. Key vibrational assignments are annotated. The transformation from 3389 cmLJ¹ (extract O-H) to 3320 cmLJ¹ (CQD O-H), disappearance of 1644 cmLJ¹, and emergence of 1044 cmLJ¹ C-O-C confirm microwave-driven etherification and surface functionalisation.

The HP-CQD spectrum (**Figure 2A**) exhibits a qualitatively transformed vibrational landscape consistent with partial carbonisation and surface functionalisation. The O-H/N-H band blue-shifts to 3320 cmO¹, indicating a change in hydrogen-bonding environment consistent with fewer intermolecular H-bonds after the conversion of large polyphenol molecules to surface-functionalised nanoparticles. The appearance of a resolved CHO/CHO stretching doublet at 2976 cmO¹ indicates the formation of aliphatic linkages. This arises from partial dehydrogenation and structural rearrangement of the precursor during carbonisation. A prominent absorption band at 1044 cmO¹ assigned to C-O-C ether stretching vibrations^29^. This observation is consistent with the dominant C-O contribution (42.67%) identified from XPS C 1s deconvolution. It suggests that microwave carbonisation promotes cross-linking of polyphenolic units via etherification, rather than complete oxygen removal. The peak at 879 cmO¹ corresponds to aromatic C-H out-of-plane bending. The band at 432 cmO¹ is attributed to C-C skeletal and C-O bending modes. These features confirm the preservation of aromatic ring structures and support the formation of a graphitic carbon core. Notably, the sharp band at 1644 cmO¹, initially attributed to isolated carbonyl groups, was no longer observed. Its redistribution into the ether and skeletal regions indicates substantial chemical transformation. This suggests that the microwave process drives polycondensation and aromatisation of the polyphenolic precursor, rather than simple oxidation. The resulting conjugated surface environment is consistent with the origin of the observed red emission.

#### 2.2.2 XPS Survey and Elemental Composition

XPS survey spectra of HP-CQDs and the parent extract (Figure 3A, B) reveal three elements in both materials: carbon (C 1s, ∼284-285 eV), oxygen (O 1s, ∼532 eV), and nitrogen (N 1s, ∼400 eV). HP-CQDs comprise C 67.24%, O 31.25%, and N 1.52%, while the extract yields C 67.41%, O 29.57%, and N 2.51%. That the carbon content remains virtually unchanged between precursor and product suggests microwave irradiation does not disturb the bulk carbon-to-heteroatom ratio; its primary effect is surface functionalisation and intramolecular reorganisation^30,31^. The slight oxygen enrichment in HP-CQDs points to selective oxidation of surface carbon atoms during aqueous microwave processing a transformation that generates the C=O and O-C=O groups widely associated with red-emitting surface states in CQDs. Nitrogen, by contrast, drops from 2.51% to 1.52%, most plausibly through volatilisation of amino-derived fragments such as ammonia under the thermal extremes of microwave treatment. The nitrogen that remains is likely embedded within the graphitic framework as pyridinic or pyrrolic species. Although present only at trace levels, such lattice nitrogen is well established as an electronic dopant in CQDs; here, it may play a role in the anomalous long-wavelength emission by narrowing the optical gap of the conjugated core.

**Figure 3.**
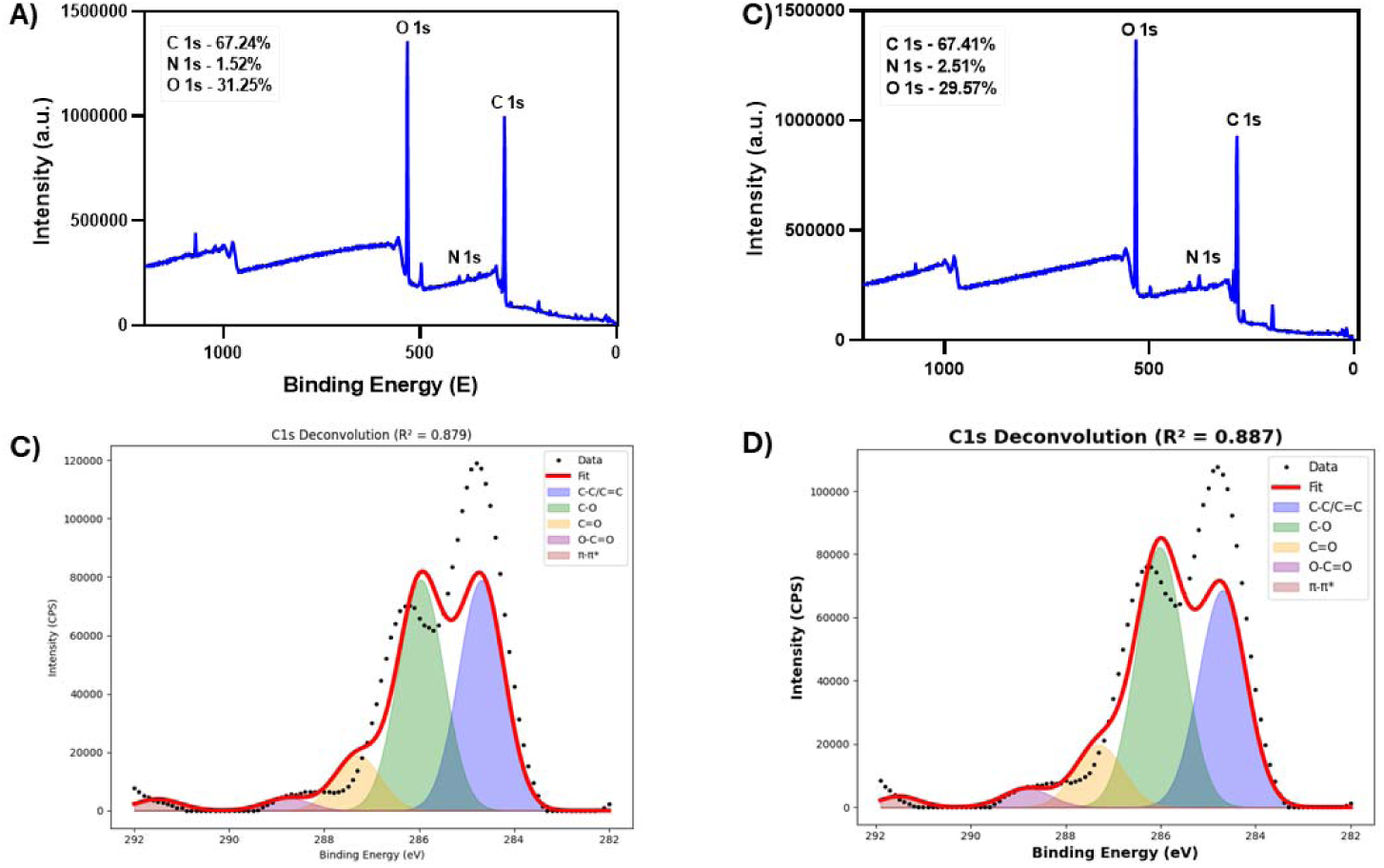
XPS characterisation. (A, B) Survey spectra of HP-CQDs and HP extract showing C 1s, O 1s, and N 1s photoelectron lines with elemental compositions annotated. (C, D) High-resolution C 1s spectra with Voigt deconvolution into C-C/C=C (284.68 eV), C-O (285.96 eV), C=O (287.30 eV), O-C=O (288.76 eV), and π-π* (291.50 eV) components for CQDs and extract, respectively. The increase in C-C/C=C and π-π* components in the CQDs compared to the extract confirms microwave-driven graphitisation.

#### 2.2.3 High-Resolution C 1s Deconvolution and Correlation with Fluorescence

High-resolution C 1s spectra of HP-CQDs and the extract were deconvoluted into five chemically distinct components (Figures 3C, D; **Tables 1 and 2**). The quality of fit is evidenced by R² values of 0.879 and 0.887 for the CQDs and extract, respectively, confirming that the five-component model adequately describes the spectral envelope.

**Table 1.** Binding energy, FWHM, and relative area of deconvoluted C 1s XPS peaks for HP-CQDs (microwave-synthesised). R² = 0.879.

| Peak | Binding Energy (eV) | FWHM (eV) | Area (CPS) | Area (%) |
| --- | --- | --- | --- | --- |
| C-C / C=C | 284.68 | 1.133 | 114,795.9 | 42.64 |
| C-O | 285.96 | 1.133 | 114,895.6 | 42.67 |
| C=O | 287.30 | 1.133 | 27,521.7 | 10.23 |
| O-C=O | 288.76 | 1.133 | 6,361.5 | 2.36 |
| $\pi \rightarrow \pi^*$ | 291.50 | 1.125 | 5,661.6 | 2.10 |

**Table 2.** Binding energy, FWHM, and relative area of deconvoluted C 1s XPS peaks for HP Extract. R² = 0.887.

| Peak | Binding Energy (eV) | FWHM (eV) | Area (CPS) | Area (%) |
| --- | --- | --- | --- | --- |
| C-C / C=C | 284.69 | 1.180 | 108,207.1 | 38.38 |
| C-O | 286.02 | 1.180 | 129,641.5 | 45.98 |
| C=O | 287.30 | 1.180 | 31,001.7 | 10.99 |
| O-C=O | 288.81 | 1.180 | 9,328.9 | 3.31 |
| $\pi \rightarrow \pi^*$ | 291.50 | 0.982 | 3,780.1 | 1.34 |

The C-C/C=C component at 284.68 eV (42.64%) represents graphitic/aromatic carbon domains and increases from 38.38% in the extract to 42.64% in the CQDs, confirming microwave-driven aromatisation and graphitisation of the aliphatic and partially unsaturated precursor carbon. The C-O component at 285.96 eV, representing ether, hydroxyl, and epoxide carbons, remains the dominant oxygenated species (42.67% in CQDs), consistent with the C-O-C ether stretch at 1044 cmO¹ in FTIR. Notably, this component is higher in the extract (45.98%), suggesting that the ether-type oxygenation is partially preserved during microwave synthesis, some C-O bonds are converted to C=O (287.30 eV, 10.23%) and O-C=O (288.76 eV, 2.36%) groups through oxidative processes.

The diagnostic π-π* shake-up satellite at 291.50 eV deserves particular mechanistic attention: its relative area increases from 1.34% in the extract to 2.10% in the CQDs, a 57% proportional increase. This satellite arises exclusively from loss processes in fully aromatic (graphitic) carbon where the photoelectron excites a π-π* transition, and its intensity is therefore a quantitative marker of graphitic sp2 domain formation^32^. The significant increase confirms that microwave carbonisation extends the aromatic conjugation length of the precursor, generating larger graphitic clusters capable of sustaining delocalised π-electron systems. This extended conjugation, combined with surface O-C=O trap states, constitutes the structural basis for the red emission: the optical gap of the graphitic core is lowered by the extended π-network, and the surface carbonyl acts as the terminal emissive state through which excited-state energy is dissipated radiatively at 675 nm.

### 2.3 Structural Characterisation

#### 2.3.1 Atomic Force Microscopy

AFM images of HP-CQDs deposited on freshly cleaved mica (**Figure 4A, B**) show uniformly dispersed nanoparticles. It appear as bright protrusions on an atomically flat background. The 2D topographic image (scale bar: 2 µm) and the corresponding 3D reconstruction both confirm the absence of significant agglomeration. This indicates good colloidal stability under the measurement conditions. The height profile reaches a maximum of ∼2.81 nm. This is consistent with sub-5 nm dimensions expected for few-layer graphitic CQDs. AFM therefore reflects the dry core particle size, without contributions from hydration layers^33^. In contrast, DLS yields a larger hydrodynamic diameter of 3.9 nm (vide infra). This difference arises from solvation effects and adsorbed water molecules. The close agreement between AFM heights (< 3 nm) and the DLS primary peak (3.9 nm) suggests that the particles lack extensive surface-bound polymer layers. Such coatings would otherwise produce a larger discrepancy between the two size measurements.

**Figure 4.**
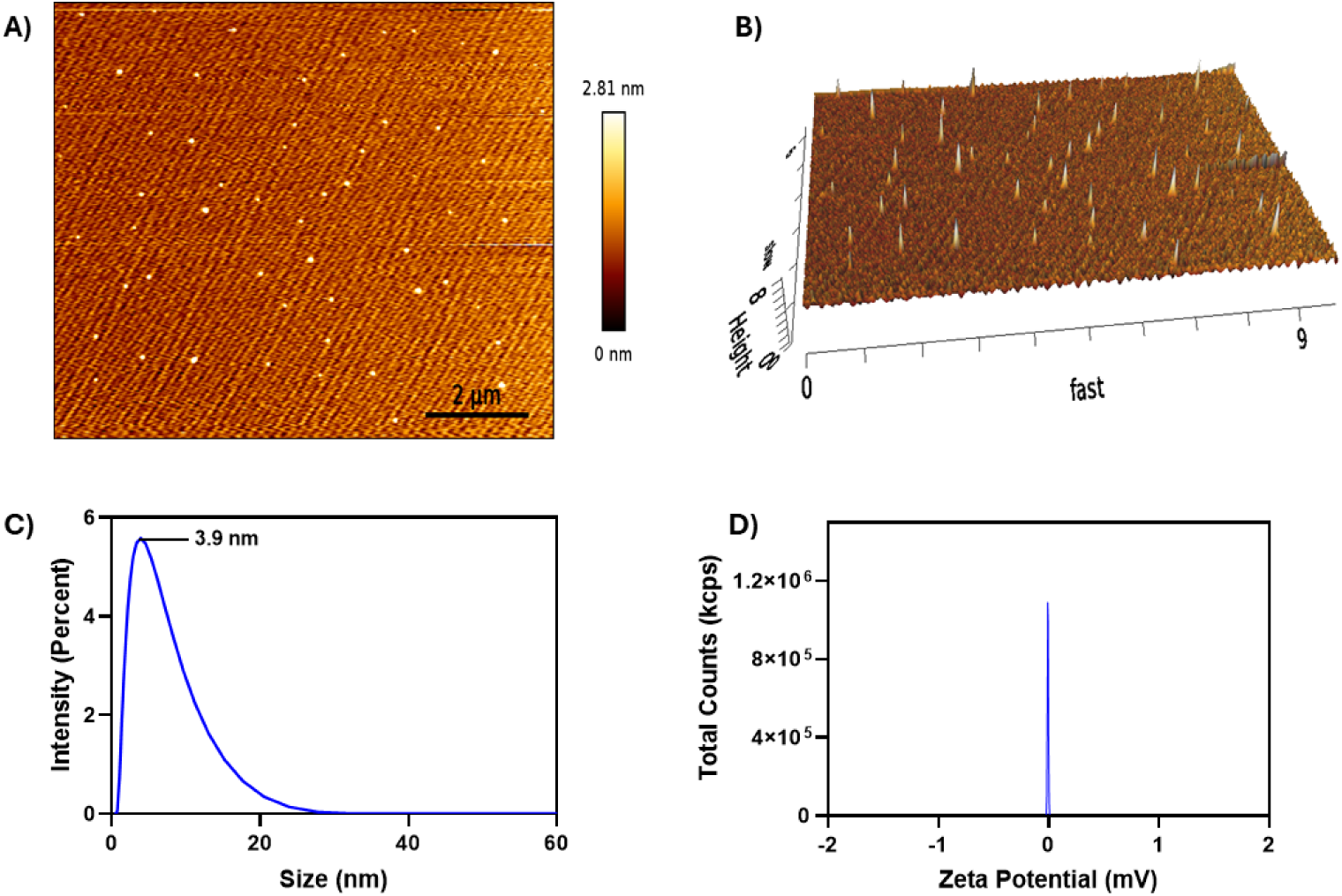
Structural characterisation of HP-CQDs. (A) 2D tapping-mode AFM image on mica substrate (scale bar: 2 µm; height scale: 0-2.81 nm) showing well-dispersed nanoparticles. (B) 3D AFM surface reconstruction confirming sub-3 nm particle heights. (C) DLS intensity-weighted size distribution with primary peak at 3.9 nm. (D) Zeta potential distribution showing near-neutral charge (∼0 mV) with high count rate.

From a biological perspective, the sub-5 nm size regime imparts several functional advantages. Such small dimensions favour efficient cellular internalisation. Uptake is typically mediated through clathrin- or caveolin-dependent endocytic pathways. Particles in this size range also exhibit improved clearance profiles^34^. Being below the ∼6 nm renal filtration threshold, they can be eliminated more readily through physiological pathways. Once internalised, their small size imposes minimal steric hindrance. This facilitates effective intracellular diffusion within the cytoplasm. These features are consistent with the efficient cellular uptake observed in confocal imaging experiments.

#### 2.3.2 Dynamic Light Scattering Size Distribution

DLS intensity distribution analysis of the HP-CQD aqueous dispersion (**Figure 4C**) reveals a narrow and monomodal peak centred at 3.9 nm. The size distribution exhibited a sharp increase with a moderate tail extending to ∼20 nm, without any secondary population in the 100–1000 nm range. This indicates that 0.22 µm syringe filtration efficiently removed larger carbonaceous aggregates and residual precursor-derived particulates. The relatively narrow distribution further suggests controlled nucleation and growth during synthesis. It is consistent with the homogeneous and rapid heating provided by microwave irradiation. The DLS-derived particle size falls well within the sub-10 nm regime. This size range is widely regarded as optimal for intracellular imaging applications. It supports efficient cytoplasmic diffusion while maintaining nuclear exclusion.

#### 2.3.3 Zeta Potential

The zeta potential profile (**Figure 4D**) exhibits a narrow and symmetric peak centred near 0 mV. The high count rate (∼1.1 × 10O kcps) indicates a monodisperse population with near-neutral surface charge. Such near-neutral values are uncommon for CQDs. Most reported systems are anionic, typically showing −20 to −40 mV due to deprotonation of surface carboxylate and hydroxyl groups at neutral pH. The near-neutral zeta potential of HP-CQDs most likely reflects a surface charge balance between opposing functionalities: negatively charged carboxylate and hydroxyl groups are counteracted by protonated amine or ammonium species derived from nitrogen-containing phytochemicals native to *Hamelia patens*. Beyond surface chemistry, experimental variables including pH, ionic strength, and particle concentration can partially screen surface charges, collectively shifting the apparent zeta potential toward neutrality. Critically, however, a near-zero value should not be interpreted as evidence of colloidal instability, as steric and solvation forces may sustain dispersion independently of electrostatic repulsion. Steric stabilisation can dominate in such systems. Surface-associated polyhydroxyl and ether groups, as indicated by FTIR, can prevent aggregation even in the absence of strong electrostatic repulsion^35^. This interpretation is supported by the narrow DLS distribution and AFM images showing well-dispersed particles. From a biological perspective, near-neutral surface charge can reduce non-specific serum protein adsorption (corona formation) and may favour passive cellular uptake mechanisms compared to highly charged surfaces that trigger rapid opsonisation.

### 2.4 In Vitro Biocompatibility and Antioxidant Activity

#### 2.4.1 MTT Cell Viability Assay

The cytotoxicity of HP-CQDs was assessed using the MTT colorimetric assay in two biologically distinct cell lines: HeLa and RPE-1. This dual-cell model was selected to enhance the translational relevance of the biocompatibility assessment^36^. Cancer-derived cells such as HeLa are generally more resistant to nanomaterial-induced stress, whereas non-tumorigenic epithelial cells like RPE-1 are metabolically active and typically more susceptible to exogenous toxicants. Demonstrating low cytotoxicity in both systems therefore provides a more rigorous evaluation of biocompatibility than single-cell-line studies and supports the potential use of HP-CQDs as bioimaging agents.

HP-CQDs maintained cell viability exceeding 95% across all tested concentrations up to 250 µg mLO¹ in both HeLa and RPE-1 lines (Figure 5A, B). One-way ANOVA with Dunnett’s post-hoc correction revealed no statistically significant differences from the vehicle control at any concentration in either cell line (all p > 0.05). The complete absence of a dose-response relationship, coupled with near-control viability even at the highest concentration tested, provides compelling evidence that HP-CQDs lack intrinsic cytotoxic activity within this range. The exceptional biocompatibility of HP-CQDs can be rationalised at the molecular level. The surface hydroxyl and ether groups provide hydrophilicity without presenting reactive electrophilic centres that could alkylate biomacromolecules. Furthermore, the polyphenol-derived surface, retaining residual hydroxyl groups, may confer intrinsic antioxidant character that mitigates oxidative stress induced by nanoparticle internalisation a point directly tested in the DPPH assay below.

**Figure 5.**
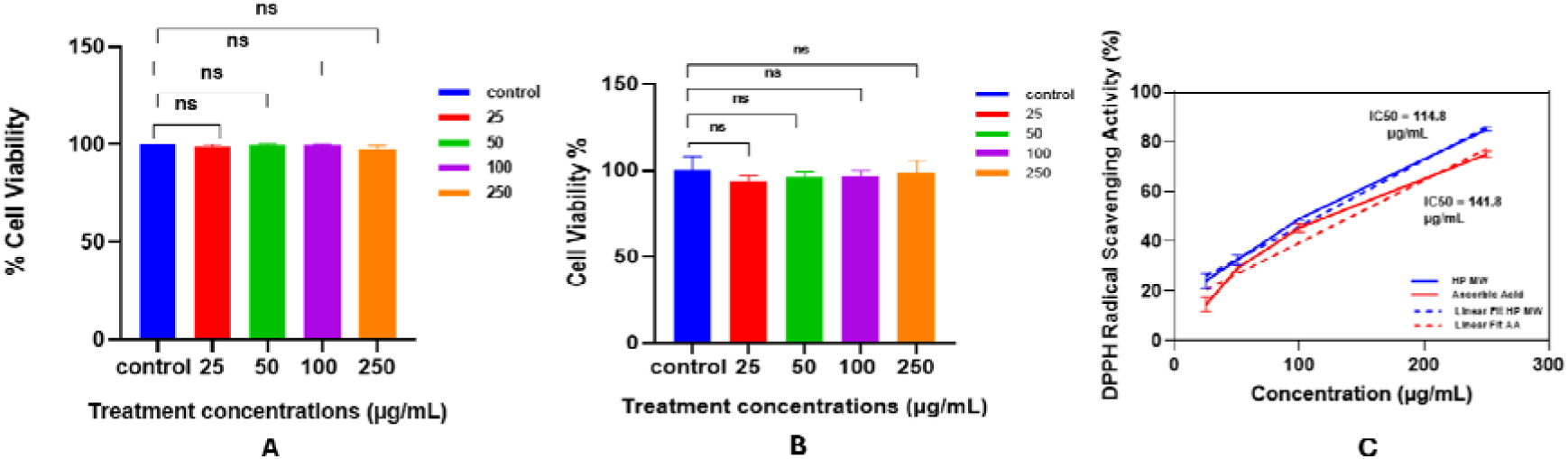
In vitro biological characterisation. (A) MTT viability assay of HP-CQDs on RPE-1 cells (25-250 µg mL[¹; n = 3, mean ± SD; one-way ANOVA, Dunnett’s test, all ns vs control). (B) MTT viability assay on HeLa cells (10–250 µg mL[¹; same statistical treatment; all ns). (C) DPPH radical scavenging activity of HP-CQDs versus ascorbic acid as a function of concentration with linear regression fits; IC[[ values are annotated (114.8 µg mL[¹ for HP-CQDs; 141.8 µg mL[¹ for ascorbic acid).

#### 2.4.2 DPPH Radical Scavenging Activity

The free radical scavenging activity of HP-CQDs was evaluated using the DPPH (2,2-diphenyl-1-picrylhydrazyl) assay, with ascorbic acid as a reference antioxidant (**Figure 5C**). Both HP-CQDs and ascorbic acid show a clear concentration-dependent scavenging response. Their dose–response profiles are closely aligned over the 20-250 µg mLO¹ range. The nearly identical slopes of the linear fits indicate comparable reaction kinetics across this concentration window. The ICOO value, defined as the concentration required to scavenge 50% of DPPH radicals, was determined by linear regression^37^. HP-CQDs exhibit an ICOO of 141.8 µg mLO¹, while ascorbic acid shows a lower value of 114.8 µg mLO¹. The ICOO ratio (141.8 / 114.8 = 1.24) indicates that HP-CQDs retain approximately 81% of the radical scavenging efficiency of ascorbic acid on a mass basis. This represents a notably high antioxidant capacity for a nanomaterial system.

This activity is mechanistically linked to the oxygen-rich surface chemistry of HP-CQDs. The high C-O content (42.67% by XPS) indicates a substantial population of hydroxyl-rich surface groups. These phenolic O-H moieties can readily donate hydrogen atoms to DPPH radicals, converting them into the stable DPPH-H form. Carbonyl functionalities (C=O, 10.23%) may further contribute through single-electron transfer pathways^38^. This provides an additional mechanism for radical neutralisation. The origin of this antioxidant behaviour is consistent with the phytochemical profile of the Hamelia patens precursor. The leaf extract contains phenolic compounds such as catechins, quercetin, and chlorogenic acid. These molecules are well known for their intrinsic radical scavenging activity^39^. Microwave carbonisation, carried out over short durations and at relatively moderate temperatures, appears to retain key functional motifs. While partial condensation of the aromatic framework occurs to form the CQD core, redox-active phenolic features are not fully eliminated. This enables preservation of antioxidant functionality in the resulting nanomaterial. The coexistence of fluorescence and antioxidant activity is particularly advantageous. HP-CQDs therefore represent a dual-functional platform. Such systems are well suited for oxidative stress sensing and theranostic applications, where real-time imaging can be coupled with simultaneous radical scavenging.

### 2.5 Cellular Uptake and Organelle Localisation in HeLa and RPE-1 Cells

Confocal laser scanning microscopy was used to evaluate the intracellular distribution of HP-CQDs in HeLa (**Figure 7**) and RPE-1 (**Figure 8**) cells. Cells were incubated for 20 min at 100, 200, and 300 µg mLLJ¹. Subcellular architecture was defined using phalloidin-FITC (green, filamentous actin cytoskeleton) and DAPI (blue, nuclear DNA). The intrinsic red fluorescence of HP-CQDs was imaged directly in the 670-720 nm detection window, without secondary labelling^8^. This strategy leverages the spectrally isolated long-wavelength emission of HP-CQDs. The signal is free from overlap with conventional organelle stains. This avoids spectral crosstalk and provides a clear advantage over blue- and green-emitting CQDs.

In HeLa cells (**Figure 7**), control samples show negligible fluorescence in the red channel. This indicates minimal cellular autofluorescence under the imaging conditions. The observed signal can therefore be attributed exclusively to internalised HP-CQDs. At 100 µg mLLJ¹, diffuse red punctate signals are discernible throughout the cytoplasm, morphologically consistent with early-stage endosomal vesicles. Quantitative fluorescence analysis (**Figure 6A, sB**) using one-way ANOVA with Dunnett’s post hoc comparison against the vehicle control shows that 100 µg mLLJ¹ yields a mean fluorescence of 112.1% of control. This corresponds to a 12.1% increase. The effect is not statistically significant (ns; p = 0.4157, q = 1.327), indicating that uptake at this dose remains below the threshold of reliable detection by whole-cell intensity analysis. At 200 µg mLLJ¹, intracellular red fluorescence increases markedly to 141.7% of control. This represents a 41.69% increase and is highly significant (p < 0.0001, q = 5.193). The increase is accompanied by a visibly denser, punctate distribution throughout the cytoplasm, consistent with enhanced intracellular accumulation.

**Figure 6.**
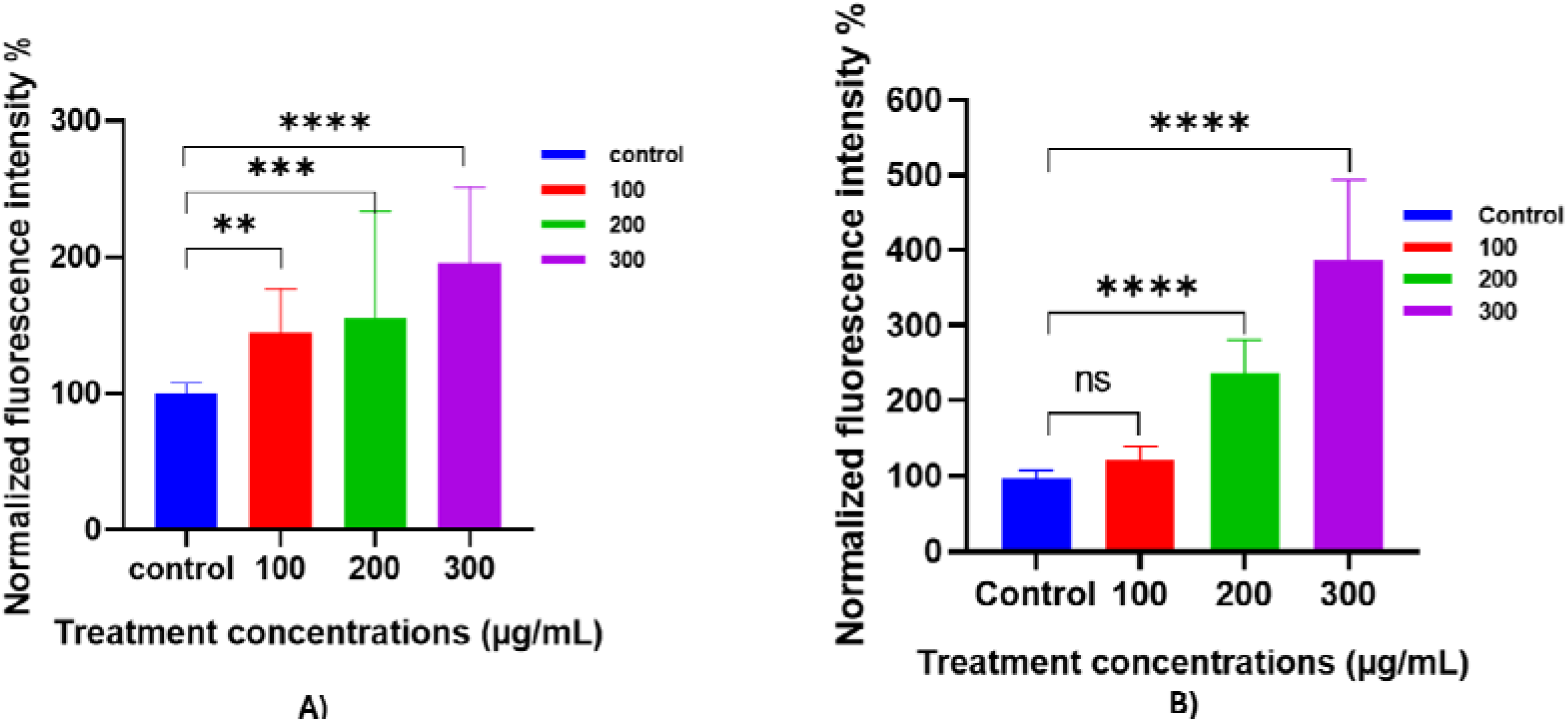
Quantitative intracellular uptake of HP-CQDs in (A) HeLa and (B) RPE-1 cells assessed by normalised red-channel fluorescence intensity (%) following 20 min incubation at 100, 200, and 300 µg mL[¹. Values are expressed as percentage of vehicle control (100%) and presented as mean ± SD (n = 30 cells per condition). Statistical analysis was performed by one-way ANOVA with Dunnett’s multiple comparisons test (α = 0.05); *** p < 0.001 vs. control; ns = not significant. RPE-1 cells exhibit markedly higher uptake efficiency than HeLa cells at equivalent concentrations, consistent with the heightened endocytic activity of retinal pigment epithelium. Significant uptake commences at 200 µg mL[¹ in both cell lines, with RPE-1 cells accumulating approximately 2.1-fold greater fluorescence intensity than HeLa cells at 300 µg mL[¹.

**Figure 7.**
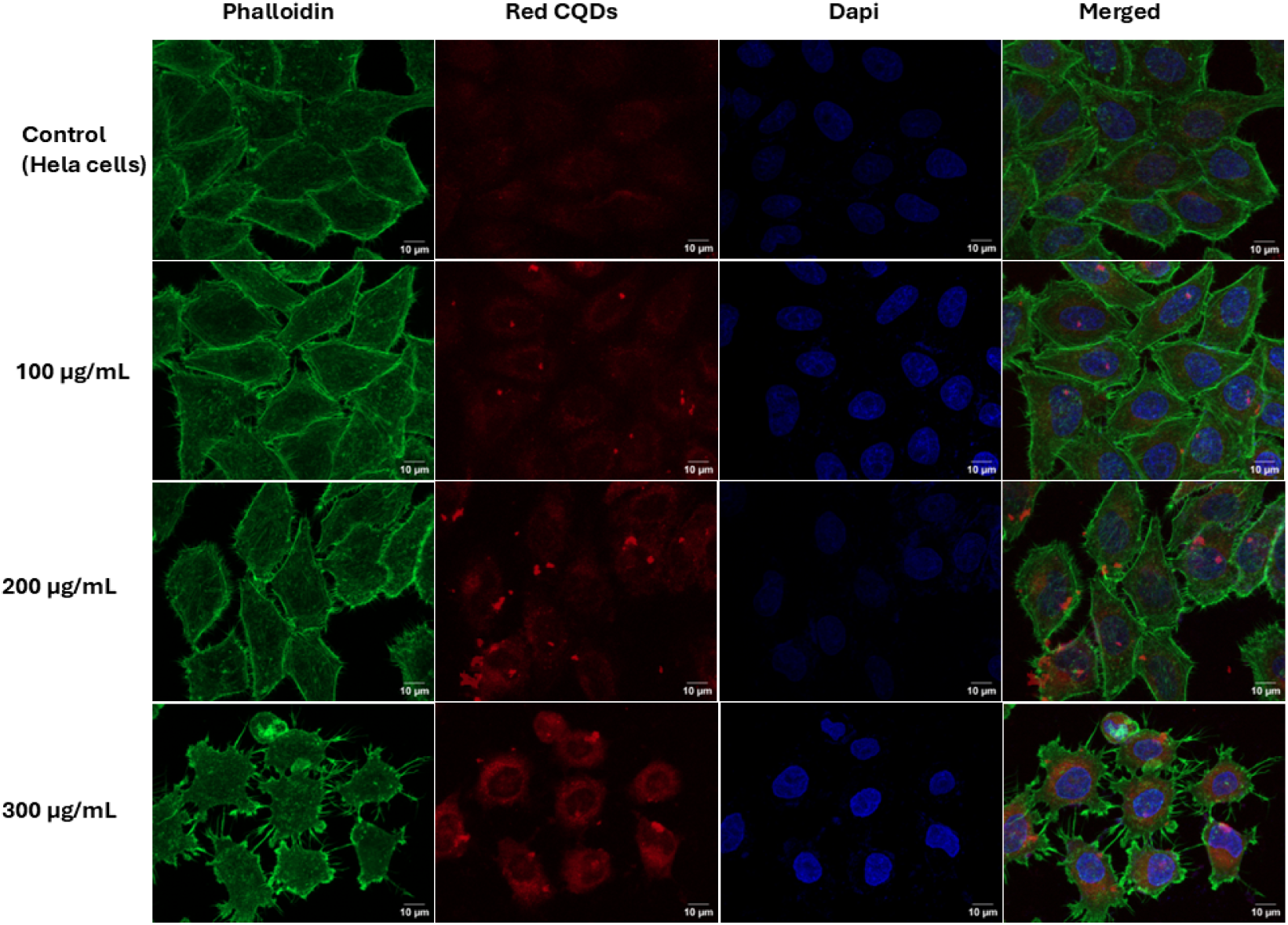
Confocal fluorescence microscopy of HeLa cells incubated with HP-CQDs for 24 h. Channels: phalloidin-FITC (green, actin), DAPI (blue, nucleus), red CQDs (red, 670–720 nm). Rows correspond to control, 100, 200, and 300 µg mL[¹. Scale bar: 10 µm. Perinuclear accumulation of red signal at 300 µg mL[¹ is indicated by the intense ring of fluorescence distinct from DAPI-stained nuclei.

**Figure 8.**
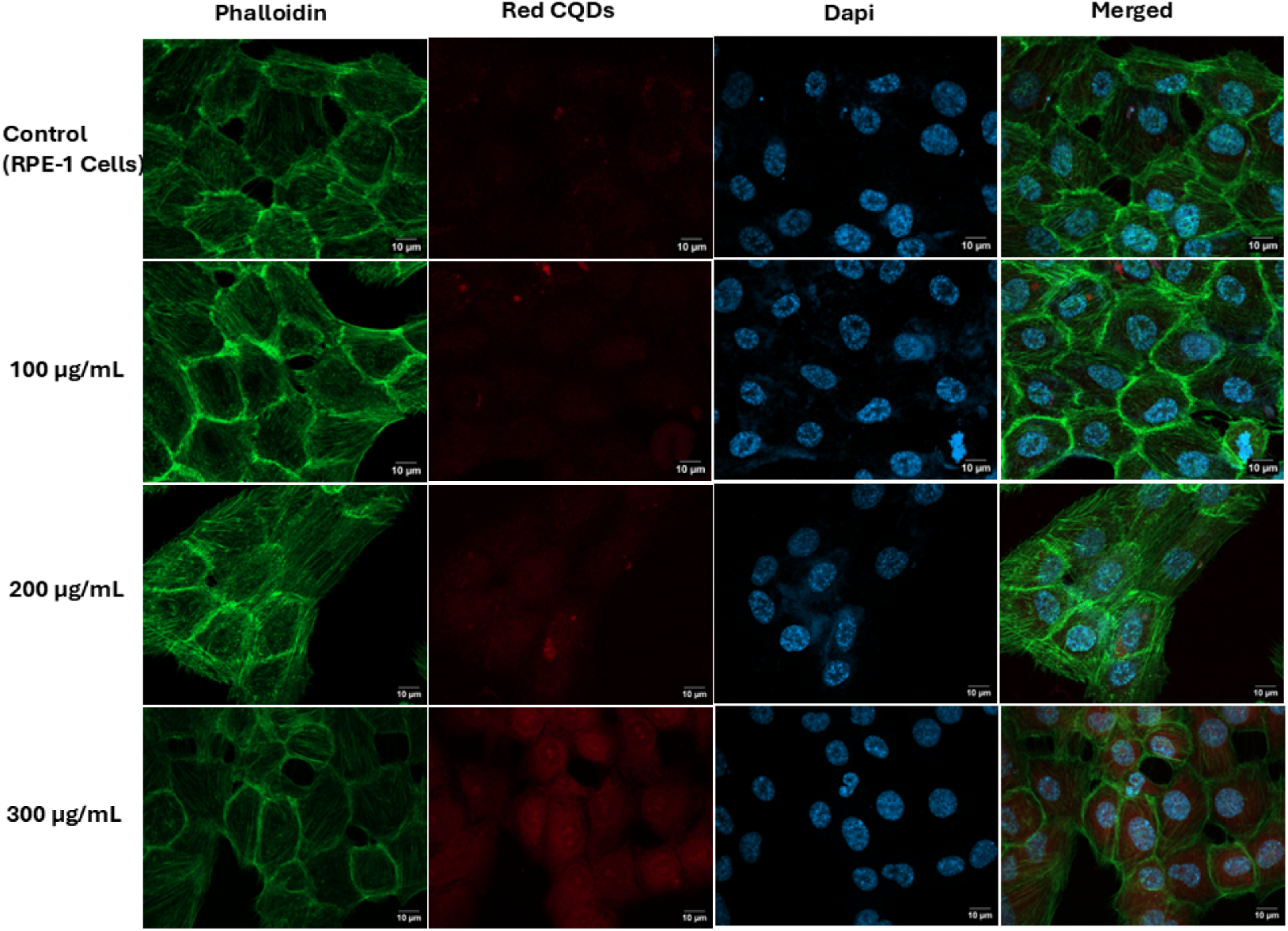
Confocal fluorescence microscopy of RPE-1 cells incubated with HP-CQDs for 24 h. Scale bar: 10 µm. Note the higher red fluorescence intensity in RPE-1 cells at equivalent concentrations compared o HeLa cells, consistent with the higher endocytic activity of retinal pigment epithelium.

The most striking and mechanistically informative observation occurs at 300 µg mLLJ¹, where the red channel reveals intense granular fluorescence organised in a perinuclear ring pattern, with mean intensity reaching 188.5% of control (mean difference 88.50%, p < 0.0001, q = 11.02). Critically, overlay of the red CQD signal with DAPI-stained nuclei in the merged images confirms complete spatial segregation of the two signals, unambiguously establishing cytoplasmic rather than nuclear localisation across all tested concentrations. RPE-1 cells (**Figure 8**) display a qualitatively similar concentration-dependent uptake trajectory, but with quantitatively superior accumulation that reflects the fundamentally distinct endocytic biology of this non-tumour epithelial line. At 100 µg mLO¹, the mean fluorescence of 121.3% versus a control of 96.24% represents a 25.06% difference that again does not reach significance (ns; p = 0.2463, q = 1.633). At 200 µg mLO¹, RPE-1 cells show a marked increase in red fluorescence, reaching 235.9% of control. This corresponds to a 139.7% increase (p < 0.0001, q = 9.108). The signal significantly exceeds that observed in HeLa cells at the same dose (141.7%). This divergence becomes more pronounced at 300 µg mLO¹. RPE-1 cells reach 385.9% of control (mean difference 289.7%, p < 0.0001, q = 18.88). This represents ∼2.1-fold higher accumulation compared to HeLa cells (188.5%) under identical conditions. The enhanced uptake in RPE-1 cells is mechanistically consistent with their physiological role. The physicochemical properties of HP-CQDs further support efficient uptake. The sub-5 nm hydrodynamic diameter (∼3.9 nm by DLS) enables compatibility with vesicle-mediated internalisation. It avoids steric constraints during membrane invagination^40^. The near-neutral surface charge reduces electrostatic repulsion from the negatively charged plasma membrane. Together, these factors facilitate efficient passive internalisation. Uptake scales with the intrinsic endocytic activity of the host cell.

At 300 µg mLO¹, both cell lines exhibit a distinct perinuclear ring of red fluorescence. The spatial localisation and morphology of this signal are consistent with accumulation in juxtanuclear compartments. These likely include late endosomes and lysosomes, which constitute the primary degradative pathway. Despite their sub-5 nm dimensions smaller than the ∼9 nm functional exclusion limit of the nuclear pore complex HP-CQDs do not enter the nucleus at any tested concentration in either cell line^41^. This indicates that nuclear entry is not governed solely by size. Instead, the particles appear to be actively excluded or preferentially retained within intracellular membrane-bound compartments. Such retention is likely favoured by interactions with anionic organelle membranes rather than passive diffusion through nuclear pores. This nuclear exclusion is functionally advantageous. Cytoplasmic and perinuclear localisation places HP-CQDs at sites of elevated reactive oxygen species (ROS) generation, including mitochondria, the endoplasmic reticulum, and lysosomal membranes^42^. Their presence in these regions aligns with their demonstrated antioxidant activity. This enables concurrent red-fluorescence imaging and localised radical scavenging within the same intracellular microenvironment.

### 2.6 In Vivo Zebrafish Biodistribution

To validate the in vivo compatibility and biodistribution capacity of HP-CQDs in a whole-organism context, zebrafish (Danio rerio) larvae at 5 days post-fertilisation (dpf) were immersed in HP-CQD solutions at 50, 100, 250, and 500 µg mLO¹ for 4 h. Brightfield, fluorescence (red channel), and merged images were acquired for each treatment group alongside vehicle-treated controls (**Figure 9a**). Normalised fluorescence intensity quantification from the merged images is shown in **Figure 9b** with statistical analysis by one-way ANOVA (Dunnett’s test vs control).

**Figure 9.**
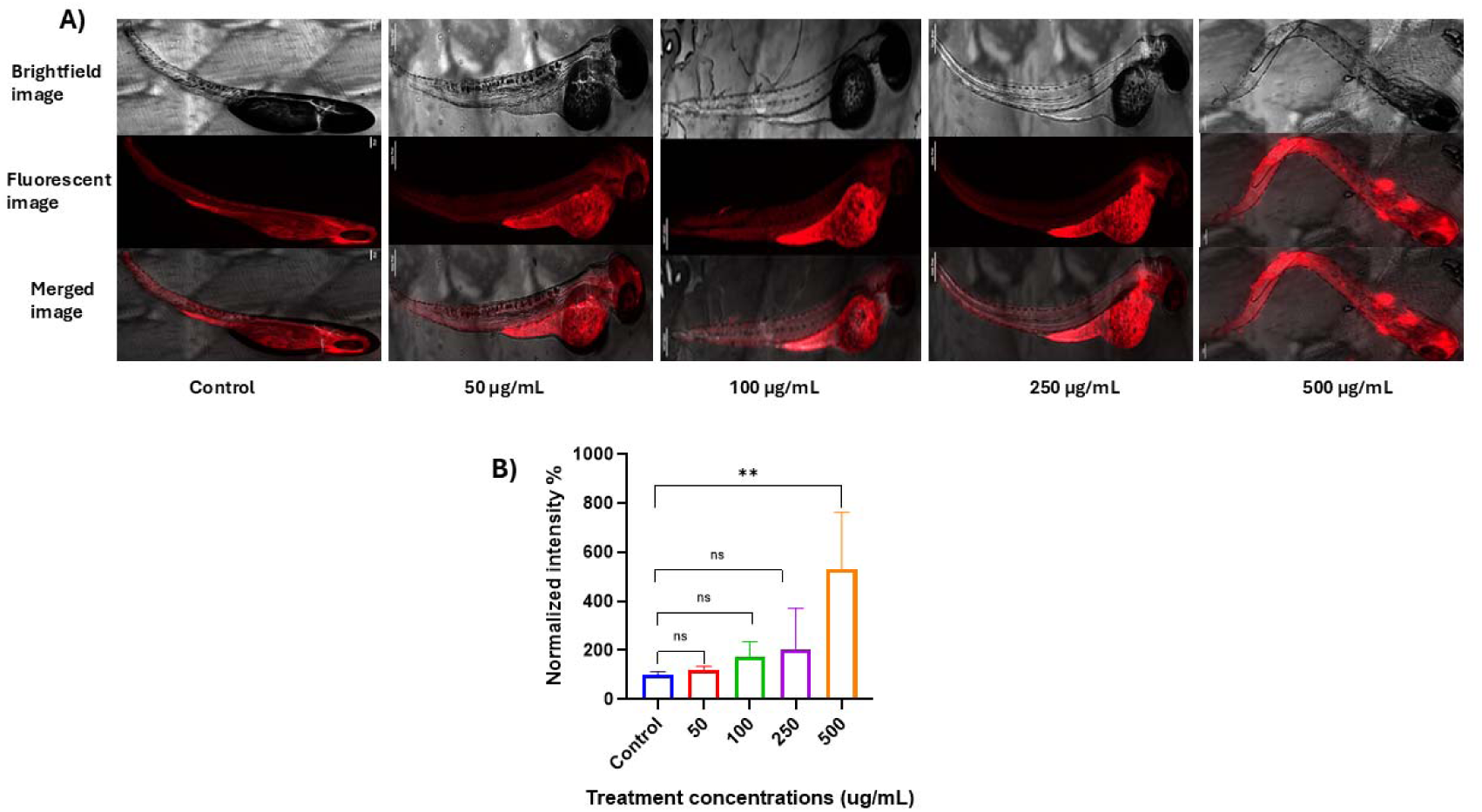
In vivo biodistribution of HP-CQDs in zebrafish larvae at 5 dpf. (A) Brightfield, fluorescence (red channel), and merged images of larvae treated with 0 (control), 50, 100, 250, and 500 µg mLLJ¹ HP-CQDs. (B) Quantification of normalised fluorescence intensity (%) across treatment groups. Data are mean ± SD; one-way ANOVA, Dunnett’s test; ** p < 0.01 vs control at 500 µg mLLJ¹; ns = not significant.

Brightfield imaging shows normal larval morphology across all treatment groups. No overt toxicity is observed up to 500 µg mLO¹. Specifically, there is no evidence of spinal curvature, yolk sac oedema, pericardial oedema, or developmental arrest. These findings provide an initial indication of in vivo biocompatibility^43^. Fluorescence imaging reveals a clear concentration-dependent increase in red signal across the larval body. At lower concentrations, fluorescence is primarily localised in the yolk sac and intestinal region. At 500 µg mLO¹, the signal extends to the trunk musculature and head region. Control larvae display negligible endogenous red fluorescence. This confirms that the observed signal originates from HP-CQD accumulation rather than intrinsic larval autofluorescence.

Quantitative analysis demonstrates that mean normalised fluorescence intensities of 113%, 175%, and 213% were observed at 50, 100, and 250 µg mLO¹, respectively (approximately estimated from the bar chart), none reaching statistical significance versus control at the experimental power level (all ns). At 500 µg mLO¹, the mean fluorescence intensity increases to ∼530% of control. This difference is statistically significant (**, p < 0.01; Dunnett’s test). These data indicate robust systemic uptake and biodistribution at this concentration. The relatively large error bars at 500 µg mLO¹ reflect inter-individual variability in larval uptake. Such variability is typical of immersion-based exposure models. It may be reduced in future studies by employing microinjection approaches to precisely control the delivered dose.

The in vivo distribution pattern, predominantly in the yolk sac, intestinal lumen, and vasculature at moderate doses is consistent with established uptake routes in zebrafish larvae. After chorion reabsorption (∼3 dpf), nanoparticles can enter via passive diffusion across the integument and through the gastrointestinal epithelium^44^. Both pathways are well documented for nanoparticle absorption. The small size of HP-CQDs (hydrodynamic diameter ∼3.9 nm) is favourable for clearance. Such particles are compatible with glomerular filtration independent pathways, including tubular secretion and hepatobiliary excretion. This reduces the likelihood of long-term bioaccumulation. These preliminary in vivo results support the suitability of HP-CQDs for whole-organism imaging. They also establish the zebrafish larval system as a practical platform for further studies. Future work can focus on dose optimisation and high-resolution, organ-specific biodistribution mapping using confocal or light-sheet microscopy.

## 3. Conclusions and Future Perspectives

This study reports the first synthesis of red-emitting carbon quantum dots from Hamelia patens leaf extract via microwave-assisted carbonisation at 750 W single-step, reagent-free process that eliminates the prolonged reaction times, toxic co-dopants, and multi-stage purification workflows that burden most long-wavelength CQD syntheses. The resulting HP-CQDs are a rare, mechanistically well-characterised example of plant-derived red fluorescence, with emission at 675 nm under 400 nm excitation spectral window where tissue autofluorescence is minimal and detector sensitivity is high.

The characterisation data presented herein supports a coherent structure–property–function narrative. UV-Vis absorption at 224 and 256 nm identifies π–π* core and n–π* surface carbonyl transitions as the primary light-harvesting processes. Excitation-wavelength-independent emission, confirmed by EEM mapping with a single dominant centre at (λem = 680 nm, λex = 401 nm), unambiguously attributes red luminescence to a discrete surface molecular state rather than a heterogeneous population of quantum-confined emitters distinction that resolves a longstanding ambiguity in CQD photophysics. XPS C 1s deconvolution substantiates this model directly: the 57% proportional increase in π–π* shake-up satellite intensity from extract to HP-CQDs quantifies microwave-driven graphitisation, while concurrent generation of surface O–C=O moieties (288.76 eV, 2.36%) and high residual C–O content (285.96 eV, 42.67%) identifies the carbonyl-functionalised graphitic surface as the structural locus of emission. ATR-FTIR corroborates this through the emergence of C–O–C ether networking (1044 cmO¹) and aliphatic linkages (2976 cmO¹), consistent with polycondensation rather than destructive pyrolysis.

AFM confirms a maximum particle height of 2.81 nm, DLS yields a narrow hydrodynamic distribution centred at 3.9 nm, and the near-neutral zeta potential (∼0 mV) points to steric rather than electrostatic colloidal stabilisation surface condition that limits non-specific biomolecular adsorption and favours biocompatibility. HP-CQDs sustain greater than 95% viability in both HeLa and RPE-1 cells up to 250 µg mLO¹, with no statistically significant cytotoxicity by Dunnett’s analysis. Beyond passive safety, HP-CQDs exhibit potent DPPH radical scavenging with an ICOO of 141.8 µg mLO¹, equivalent to roughly 81% of the antioxidant potency of ascorbic acid on a mass basis. This activity is mechanistically rooted in the high surface hydroxyl density preserved from the parent polyphenol precursor through carbonisation. The resulting dual identity fluorescent probe and antioxidant agent within a single nanomaterial distinguishes HP-CQDs from both conventional fluorescent dyes and single-function CQD systems.

Cellular uptake studies showed concentration-dependent cytoplasmic accumulation in both cell lines. Significant internalisation was observed from 200 µg mLO¹ onward (p < 0.0001). At 300 µg mLO¹, RPE-1 cells exhibited approximately 2.1-fold higher fluorescence intensity than HeLa cells (385.9% vs. 188.5% of control), consistent with the greater endocytic activity of retinal pigment epithelial cells. Despite particle sizes below 5 nm, nuclear exclusion was maintained at all concentrations. At the highest dose, fluorescence accumulated in the perinuclear region, localising HP-CQDs near intracellular sites associated with elevated oxidative stress. This distribution may be advantageous for antioxidant delivery. In vivo studies in zebrafish larvae confirmed systemic uptake, with significant fluorescence enhancement at 500 µg mLO¹ (p < 0.01) and no detectable morphological toxicity. These findings support the in vitro biocompatibility results and establish the zebrafish model as a suitable platform for future pharmacokinetic studies.

This work advances the green CQD field on five distinct fronts: (i) first exploitation of Hamelia patens as a carbon precursor, leveraging its distinctive polyphenolic and flavonoid richness to achieve red emission without synthetic reagents. (ii) mechanistic resolution of surface-state-governed red luminescence through quantitative XPS π-π* satellite analysis. (iii) demonstration that microwave carbonisation of a polyphenol-rich precursor achieves extended graphitic conjugation sufficient for long-wavelength emission within minutes. (iv) establishment of dual imaging antioxidant function from a structurally unified, single-component nanomaterial. (v) cell-line-specific quantification of uptake efficiency that reveals the RPE-1 retinal line as a particularly receptive target for HP-CQD-based intracellular applications.

Several research directions follow naturally from the findings reported here. The absolute photoluminescence quantum yield (PLQY) of HP-CQDs remains to be determined; since red-emitting CQDs in the literature span a broad range (1–35%), integrating sphere measurements would precisely position HP-CQDs within this landscape and inform any surface passivation strategies aimed at improving radiative efficiency^45^. Complementing the XPS evidence for graphitisation, HRTEM with lattice fringe analysis would directly resolve graphitic core domain size and d-spacing (∼0.21 nm for sp² carbon), enabling a more rigorous correlation between core crystallinity and emission wavelength.

The intracellular compartment targeted at high CQD doses tentatively assigned to the juxtanuclear lysosomal endosomal system based on perinuclear localisation requires confirmation through co-localisation experiments using organelle-specific fluorescent probes (LysoTracker, ER-Tracker, MitoTracker) and Pearson correlation coefficient analysis. The antioxidant activity demonstrated in the DPPH assay must similarly be validated in a cellular context. DCFH-DA fluorescence assays in HOOO-challenged cells are needed to verify intracellular ROS-scavenging activity^46^. These experiments would determine whether hydroxyl-mediated antioxidant capacity is retained in the intracellular reducing environment. Such validation is essential before advancing theranostic claims.

Systematic optimisation of microwave power, irradiation time, and solvent polarity could establish a rational synthesis parameter map. This may enable tunable emission across the red-to-near-infrared region using the same Hamelia patens precursor. Because the synthesis is single-step and reagent-free, lifecycle assessment (LCA) studies should also be performed. Comparing HP-CQD production with conventional hydrothermal methods would quantify the environmental benefits of the green synthesis approach. Such metrics are increasingly important for responsible nanomaterial development.

Collectively, these directions reframe the present work not as a terminal characterisation study but as the foundation for a broader research programme centred on red-fluorescent, antioxidant-active, biocompatible nanomaterials with genuine translational potential in retinal biology, oxidative stress theranostics, and sustainable nanophotonics.

## 4. Experimental Section

### 4.1 Materials

Fresh leaves of Hamelia patens were collected from IIT Gandhinagar Campus, in Winter season, washed thoroughly with deionised water, dried at 60 °C for 48 h, and ground into a fine powder using a mortar and pestle. Deionised water (Milli-Q) was used throughout. DPPH (2,2-diphenyl-1-picrylhydrazyl), ascorbic acid, MTT reagent, dimethyl sulfoxide (DMSO), phalloidin-FITC, and DAPI were purchased from Sigma-Aldrich and used as received.

### 4.2 Synthesis of HP-CQDs

Leaf powder of Hamelia patens (1 g) was dispersed in 10 mL of 50% (v/v) ethanol-water (1:10 w/v). The mixture was stirred magnetically for 4 h at room temperature to obtain the extract. The suspension was centrifuged at 6000 rpm for 10 min. The supernatant was collected, and insoluble plant debris was discarded. The extract was transferred to a glass vessel and subjected to microwave irradiation at 750 W. Heating was performed for 5-6 min using 30 s cycles until a dark-brown carbonaceous product formed. The crude product was redispersed in 10 mL ethanol-water. It was filtered sequentially through Whatman No. 1 paper and a 0.22 µm PTFE syringe filter to remove residual carbon aggregates. The filtrate was concentrated by rotary evaporation at 60 °C under reduced pressure. The residue was redissolved in 5 mL deionised water and lyophilised for 48 h to yield HP-CQDs as a dry powder. The final product was stored at 4 °C in the dark until further use.

### 4.3 Characterisation

UV-Vis absorption spectra were recorded on Thermo Scientific Evolution Pro UV-Vis Spectrophotometer from 200 to 800 nm. Steady-state photoluminescence spectra and excitation-dependent PL spectra (λex = 250-500 nm, step 50 nm) were acquired using a HORIBA Fluorolog spectrofluorometer. Three-dimensional excitation-emission matrix (EEM) spectra were collected over the same excitation range with emission recorded from 600 to 800 nm. ATR-FTIR spectra were recorded using a PerkinElmer Frontier FT-IR equipped with a diamond ATR crystal over the spectral range 4000-400 cmO¹ at a resolution of 4 cmO¹ with 64 accumulated scans. XPS measurements were performed on a Thermo Scientific K-Alpha. Survey scans were acquired at 160 eV pass energy; high-resolution C 1s spectra at 20 eV pass energy. XPS C1s spectra were deconvoluted using Gaussian peak functions in the lmfit package after iterative polynomial background subtraction. Peak positions were constrained within chemically reasonable limits, and selected component FWHM values were linked during fitting to maintain spectral consistency. DLS and zeta potential measurements were performed on a Malvern Zetasizer ultra at 25 °C. Tapping-mode AFM images were acquired on a High-Performance Biological AFM at room temperature; samples were prepared by depositing dilute CQD dispersion (0.1 mg mLO¹) on freshly cleaved mica and vacuum-drying in desiccator.

### 4.4 MTT Cell Viability Assay

HeLa and RPE-1 cells were cultured in DMEM supplemented with 10% fetal bovine serum (FBS) and 1% penicillin-streptomycin at 37 °C in 5% COO. Cells were seeded in 96-well plates (5 × 10³ cells/well) and allowed to adhere for 24 h. HP-CQDs (dissolved in culture medium to final concentrations of 10, 25, 50, 100, and 250 µg mLO¹) were added and incubated for 24 h. MTT reagent (0.5 mg mLO¹ final concentration) was added and incubated for 4 h. Formazan crystals were dissolved in DMSO (100 µL/well) and absorbance measured at 570 nm. Cell viability (%) was calculated relative to vehicle control (100%). Data are presented as mean ± SD (n = 4); statistical analysis by one-way ANOVA with Dunnett’s multiple comparisons test (α = 0.05) using GraphPad Prism.

### 4.5 DPPH Radical Scavenging Assay

DPPH (0.1 mM in ethanol) was mixed with HP-CQD solutions at concentrations of 10-250 µg mLO¹ in a 1:1 v/v ratio. After 30 min incubation at room temperature in the dark, absorbance at 517 nm was measured. Scavenging activity (%) was calculated as [(AO - AO) / AO] × 100, where AO is the absorbance of DPPH without sample and AO is the absorbance with sample. ICOO values were determined by linear regression of scavenging activity (%) versus concentration. Ascorbic acid was used as positive control. Data are mean ± SD, n = 3.

### 4.6 Confocal Fluorescence Microscopy

HeLa and RPE-1 cells were seeded onto 35 mm glass-bottom dishes at a density of 1 × 10O cells per dish and allowed to adhere for 24 h. HP-CQDs dispersed in complete culture medium (100, 200, and 300 µg mLO¹) were then added, and cells were incubated for 20 min at 37 °C under a humidified atmosphere containing 5% COO. Following incubation, cells were fixed with 4% paraformaldehyde for 15 min and permeabilised using 0.1% Triton X-100 for an additional 15 min. After PBS washing, the actin cytoskeleton and nuclei were stained with phalloidin-FITC (1:500 dilution; ex/em: 488/519 nm) and DAPI (1 µg mLO¹; ex/em: 405/461 nm), respectively. HP-CQDs were visualised directly in the red fluorescence channel using excitation at 405 or 488 nm and emission collection between 670–720 nm. Confocal fluorescence images were acquired using a Leica TCS SP8 fitted with a 63× oil-immersion objective lens (NA 1.4). Fluorescence intensity analysis was performed in ImageJ (NIH) using 30 cells per condition, and statistical significance was evaluated using Dunnett’s multiple-comparison test.

### 4.7 Zebrafish Biodistribution

Zebrafish (Danio rerio) maintenance and experimental procedures were performed in accordance with the guidelines approved by the Institutional Animal Ethics Committee. Zebrafish larvae at 5 days post-fertilisation (dpf) were transferred to 6-well plates containing E3 embryo medium, with five larvae per well. HP-CQDs dispersed in E3 medium at concentrations of 50, 100, 250, and 500 µg mLO¹ were added, and the larvae were incubated for 4 h at 28 °C in the dark. Following exposure, larvae were washed thoroughly with fresh E3 medium and fixed using 4% paraformaldehyde (PFA). Brightfield and fluorescence images in the red emission channel were acquired using a Leica TCS SP8. Fluorescence intensities were quantified relative to untreated control larvae and analysed using one-way ANOVA followed by Dunnett’s multiple-comparison test, with at least five larvae analysed per experimental group.

## Declarations

None

## Acknowledgements

The authors express their gratitude to every member of the D.B. study group for their helpful criticism and thorough evaluation of the work. We sincerely thank the Indian Institute of Technology Gandhinagar for its financial and infrastructure support. The Central Instrumentation Facility (CIF) at IIT Gandhinagar is acknowledged by the authors for granting access to the characterisation tools utilised in this investigation, including AFM, FTIR, XRD. A.S. thanks financial support from the Prime Minister’s Research Fellowship and the Government of India’s Ministry of Education. The Indian National Young Academy of Sciences acknowledges D.B. as a member. J.P and S.B. thanks IIT Gandhinagar’s Department of Biological Sciences and Engineering for providing the resources and research environment needed to complete this work. The work in the host labs is supported by funding from ANRF-CRG, GSBTM, MoES-STARS, CCRH, IITGN.

## Author Contributions

D.B.: Funding acquisition, writing, review and editing, supervision, resources, and conceptualisation. S.B, J.P., B.P., A.S, G.P. S.J: Conceptualisation, research, formal analysis, methodology, first draft writing, and visualisation. S.B.: Research, Approach. AKM and A.S.:

Data curation, methodology, and investigation.

